# Cryo-EM structure of a proton-activated chloride channel TMEM206

**DOI:** 10.1101/2020.11.05.369835

**Authors:** Zengqin Deng, Yonghui Zhao, Jing Feng, Jingying Zhang, Haiyan Zhao, Michael J. Rau, James A.J. Fitzpatrick, Hongzhen Hu, Peng Yuan

## Abstract

TMEM206 has been recently identified as an evolutionarily conserved chloride channel that underlies ubiquitously expressed, proton-activated, outwardly rectifying anion currents. Here we report the cryo-electron microscopy structure of pufferfish TMEM206, which forms a trimeric channel, with each subunit comprising two transmembrane segments, the outer and inner helices, and a large extracellular domain. An ample vestibule in the extracellular region is accessible laterally from the three side portals. The central pore contains multiple constrictions preventing ion conduction. A conserved lysine residue near the cytoplasmic end of the inner helix forms the presumed chloride ion selectivity filter. Unprecedentedly, the core structure and assembly closely resemble those of the epithelial sodium channel/degenerin family of sodium channels that are unrelated in amino acid sequence and conduct cations instead of anions. Together with electrophysiology, this work provides insights into ion conduction and gating for a new class of chloride channels that is architecturally distinct from previously characterized chloride channel families.

## Introduction

Chloride ions (Cl^−^) are the most abundant anions in animals, thus Cl^−^ movement across cell membranes, mediated by Cl^−^ channels and transporters, is central to numerous cellular functions, such as regulation of cell volume, acidification of intracellular vesicles, and excitability control in muscle cells (*1*, *2*). Widely observed in mammalian cells, the acid- or proton-activated outwardly-rectifying Cl^−^ currents (*I*_Cl,H_, also referred to as ASOR or PAORAC) have long been recognized but the molecular components behind these currents have remained elusive until very recently (*3*–*12*). Two independent studies using genome-wide RNA interference screen have identified TMEM206 as the underlying anion channel (*13*, *14*). Evolutionally conserved in vertebrates, TMEM206 recapitulates biophysical characteristics of the proton-activated Cl^−^ currents, including an outwardly-rectifying current-voltage (I–V) relationship and a permeability sequence of SCN^−^ > I^−^ > NO_3_^−^ > Br^−^ > Cl^−^ (*13*, *14*). The presence of *I*_Cl,H_ in all mammalian cell types examined so far suggests that TMEM206 may be universally expressed in all tissues and play an essential role in cellular responses to extracellular acidification. Interestingly though, the channel is activated at a highly acidic extracellular pH (~5.5-6.0) that may be limited to pathological conditions, such as cancer and ischemic stroke (*9*, *13*–*18*) in which the *I*_Cl,H_ currents may contribute to cell death induced by tissue acidosis (*6*, *19*). Consistent with this notion, deletion of TMEM206 attenuated acid-induced cell death (*13*, *14*), suggesting that pharmacological inhibition of TMEM206 potentially alleviates acidosis-associated pathologies.

Cl^−^ channels are remarkably diverse in both amino acid sequence and three-dimensional architecture, as demonstrated by the structurally and functionally characterized Cl^−^ channel families, including CLC channels (*1*), Bestrophin (*20*–*22*), TMEM16 (*23*–*26*), cystic fibrosis transmembrane conductance regulator (CFTR) (*27*), anion-selective Cys-loop receptors (*28*), and volume-regulated anion channels LRRC8/VRAC (*29*–*31*). The lack of signature sequences or structural motifs among these channels has posed significant challenges in regard to the molecular identification of Cl^−^ currents, as is the case of TMEM206 (*13*, *14*). Without discernible sequence homology with previously characterized Cl^−^ channels, TMEM206 likely represents a new class of Cl^−^ channels with distinct subunit architecture, channel assembly, and ion conduction and activation mechanisms. Here we present a cryo-electron microscopy (cryo-EM) structure of pufferfish TMEM206, which reveals a trimeric channel architecture that is indeed different from previously known Cl^−^ channels. In combination with electrophysiology, this work provides the first structural and functional description of an evolutionarily conserved and broadly expressed Cl^−^ channel and establishes a molecular framework for understanding Cl^−^ conduction and channel gating.

## Results

### Fusion strategy for cryo-EM structure determination

To identify TMEM206 candidates for structural studies, we expressed multiple orthologs with a C-terminal green-fluorescent protein (GFP) in yeast *Pichia pastoris* and analyzed channel expression and assembly profiles using fluorescence-detection size-exclusion chromatography (FSEC) (*32*). Subsequent large-scale purification identified pufferfish (*Takifugu rubripes*) TMEM206, which shares 50% sequence identity to the human channel (fig. S1), as a promising target indicated by a well-resolved oligomeric assembly on size-exclusion chromatography (fig. S2). The full-length wild-type pufferfish TMEM206 protein was purified to homogeneity and subjected to single-particle cryo-EM analysis. Three-dimensional reconstruction yielded a low resolution (~6.2 Å) map that revealed a trimeric channel architecture with both transmembrane and extramembrane domains (fig. S3). The trimeric channel has a predicted molecular weight of ~120 kDa and likely contains unstructured segments. Thus, the relatively small particle size may present a challenge in high-resolution cryo-EM reconstruction. Further, particles were sparsely distributed on the cryo-EM grids (fig. S2), and preparations of grids at higher protein concentrations failed to improve the particle density because channels were increasingly inclined to aggregate upon vitrification.

To overcome these technical difficulties, we fused the channel with a C-terminal BRIL (thermo-stabilized apocytochrome b_562_RIL), a four-helix bundle protein that has been widely used as a crystallization chaperone to improve membrane protein stability and to promote crystal formation (*33*, *34*). To potentially facilitate particle alignment with the additional molecular mass from BRIL, we systematically shortened the extreme C-terminus of pufferfish TMEM206, which is not conserved among orthologs and is most likely unstructured, to increase the overall structural rigidity between the channel and BRIL. On the basis of FSEC profiles, we selected a construct with the last four C-terminal amino acids removed. This construct, we termed TMEM206_EM_, contains amino acids 1-349 of pufferfish TMEM206 and BRIL. By contrast, purified TMEM206_EM_ showed markedly reduced aggregation on size-exclusion chromatography and produced densely distributed particles on cryo-EM grids (fig. S2). These improvements allowed us to obtain a cryo-EM reconstruction with an overall resolution of ~3.5 Å with C3 symmetry imposed (fig. S4 and S5, Table S1). The quality of the cryo-EM density map is sufficient for *de novo* model building guided by bulky side chains. Portions of the N- and C-termini (amino acids 1-64 and 335-349) and BRIL were not resolved in the density map and thus were excluded from the model. The final atomic model, consisting of residues 65-159, 168-251 and 255-334, has good stereochemistry and fits well into the density (fig. S5 and Table S1). The model also matches the lower resolution map calculated from the intact wild-type channel, indicating that the BRIL fusion did not undermine the structural integrity of the channel (fig. S6). Furthermore, when expressed in *TMEM206*-knockout HEK293T cells, pufferfish TMEM206 and TMEM206_EM_ displayed a similar I-V relationship, pH dose response, and anion selectivity (Fig. 1, A-D, fig. S7 and S8).

**Figure 1.**
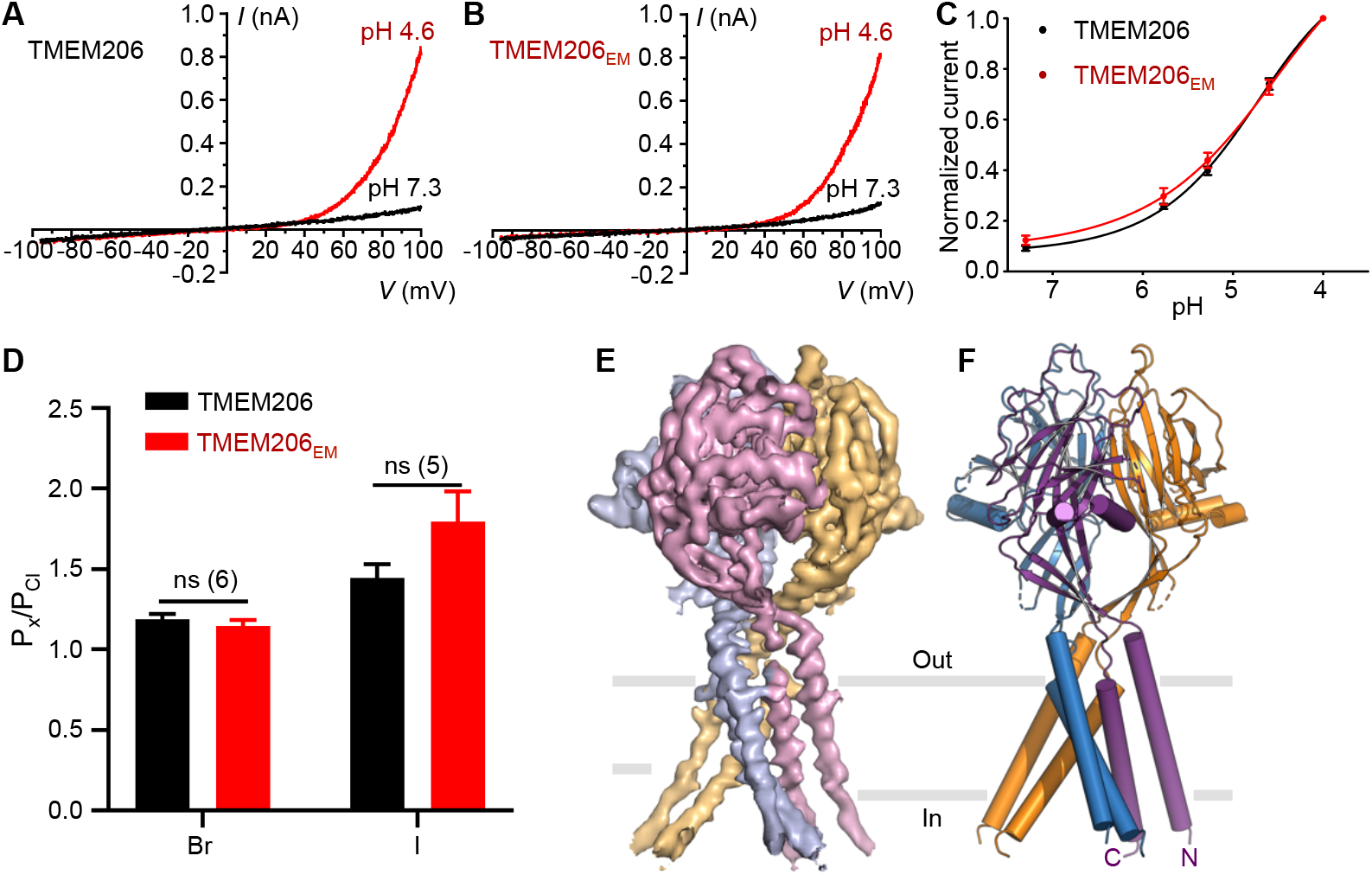
Function and structure of pufferfish TMEM206. (**A**-**B**) Representative whole-cell current traces activated by extracellular pH 4.6 for pufferfish TMEM206 (**A**) and TMEM206_EM_ (**B**). Channel constructs were expressed in *TMEM206* knockout HEK293T cells. (**C**) Normalized current-to-pH relationships of pufferfish TMEM206 (n = 6-9 cells per data point) and TMEM206_EM_ (n = 5-6 cells per data point). All currents were recorded at room temperature and normalized to pH 4.0 currents at +100 mV. (**D**) Anion selectivity for pufferfish TMEM206 and TMEM206_EM_. Data are presented as mean ± SEM (n.s, not significant, Student’s t-test). (**E**) Cryo-EM density of pufferfish TMEM206_EM_ contoured at 7.0 σ and colored by individual subunits. (**F**) Trimeric structure of pufferfish TMEM206_EM_.

### A trimeric Cl^−^channel

TMEM206 forms a symmetric trimer, with each subunit containing a transmembrane domain (TMD) with two membrane-spanning helices (TM1 and TM2) and a large extracellular domain (ECD) enriched in β strands (Fig. 1, E and F). The outer helix TM1 and inner helix TM2 within a single subunit are arranged in an approximately antiparallel fashion, tilted by ~30° to the membrane normal. The ECD comprises an inner β-domain arranged around the central three-fold symmetry axis and an outer β-domain with a helix-turn-helix (HTH) insertion positioned at the periphery of the channel (Fig. 2, A-C). The inner β-domain, consisting of β1, β3, β6 and β9-12, is further organized into the upper and lower layers that are held together by an elongated pair of antiparallel β-strands β9 and β10. β1 and β12 in the lower layer are connected to TM1 and TM2, respectively. Besides the β9-β10 pair, the upper layer contains three additional strands, β3, β6 and β11, and connects to the lower layer via a short β11-β12 linker. In contrast to a compact lower layer, the upper layer of the ECD is expanded by the peripheral outer β-domain, consisting of β2, β4, β5, β7 and β8, and the HTH insertion between β7 and β8. Extensive side-chain contacts, mainly through van der Waals interactions, are involved in the interface between the inner and outer β-domains.

**Figure 2.**
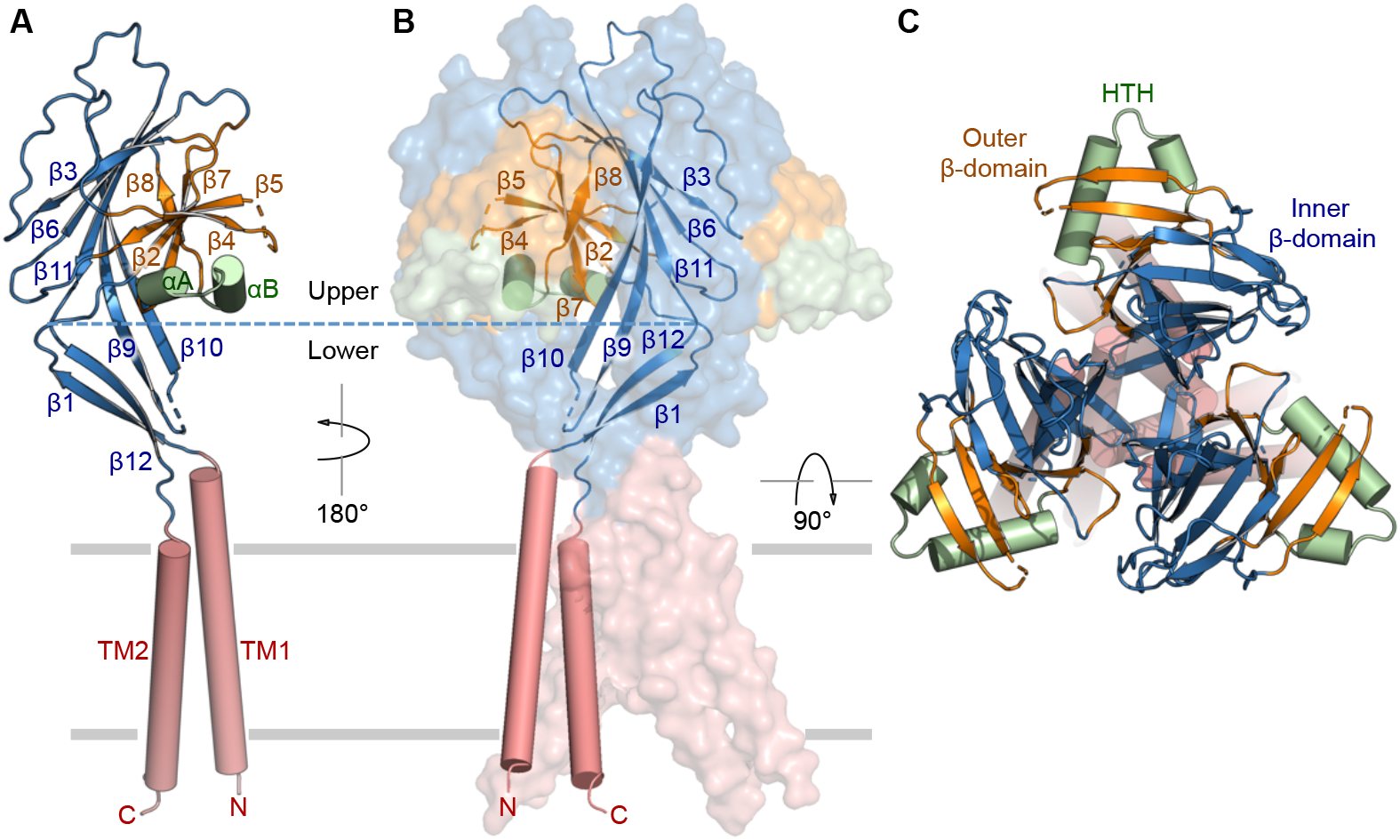
Subunit structure and channel assembly. (**A**) Structure of a single subunit, showing the transmembrane domain (red), inner β-domain (blue), outer β-domain (orange), and helix-turn-helix (green). Secondary structure elements are indicated. (**B**) The trimeric channel assembly. Two of the subunits are shown in surface representation. (**C**) Orthogonal view as in (**B**), from the extracellular side.

The inter-subunit interface, contributed by the ECD and TMD, buries ~2,400 Å^2^ of molecular surface per subunit. In the extracellular region, intimate packing interactions are limited to two regions, the top portion of the ECD and the ECD-TMD junction, leaving considerable empty spaces in the middle between subunits (Fig. 3). The three subunits come to close proximity at the very top of the ECD, with side chains of residues F238 and K267 facing the central three-fold symmetry axis (Fig. 3A). On the side, loops from neighboring subunits interdigitate through a network of both van der Waals and hydrogen bonding interactions (Fig. 3B). In particular, the aromatic side chain of F198 is nestled in the hydrophobic pocket from an adjacent subunit composed of several aromatic side chains from F186, F268 and F283, and is within the distance of cation-π interaction with R239 from the neighboring subunit. The robust inter-subunit interactions at the uppermost portion of the ECD may contribute to a stationary structural scaffold that supports gating transitions necessary at the distal TMD and ECD-TMD junction.

**Figure 3.**
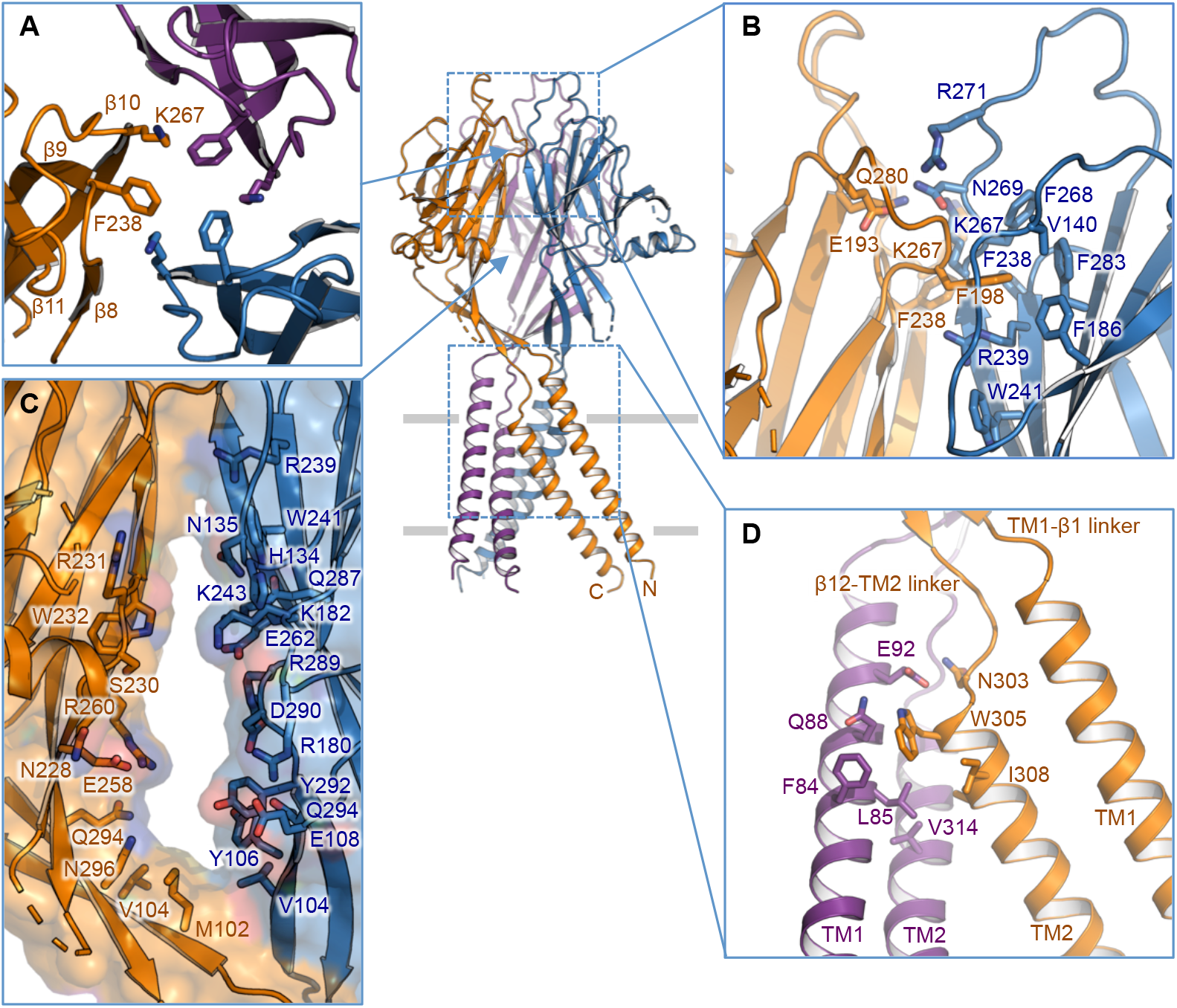
Inter-subunit interface. (**A**) The trimeric interface at the apex of the extracellular domain (ECD). Side chains of K267 and F238 are highlighted. (**B**) Side view of the inter-subunit interface at the top layer of the ECD. Residues involved in the interface are shown in stick representation. (**C**) The side portal in the middle of the ECD between two neighboring subunits. Surface and residues lining the wall are illustrated. (**D**) The TM1-TM2 inter-subunit interface.

The trimeric channel assembly introduces three lateral openings (side portals) in the middle of the extracellular region (Fig. 3C). In each portal, the interior wall is predominantly lined by polar and charged side chains, likely facilitating ion and water passage. The elongated side portals extend to the ECD-TMD junction where tight packing interactions resume. The short linkers of TM1-β1 and β12-TM2 create a narrow ‘neck’ immediately above the lipid membrane (Fig. 3D). Within the membrane, the trimer interface is primarily mediated by the inner helix TM2, which, together with the ‘neck’, restricts the central ion-conduction path. Side chain interactions are also observed between TM2 and an adjacent TM1 on the extracellular end (Fig. 3D). Notably, this interface reveals the close proximity between L85 in TM1 and W305 in TM2. Consistent with the notion that the TM1-TM2 inter-subunit interface is likely involved in regulation of channel activity, cysteine substitutions at the equivalent positions of L85 and W305 in human TMEM206 both increased *I*_Cl,H_ currents at negative voltages with the application of cysteine-modifying reagent MTSES (2-Sulfonatoethyl methanethiosulfonate) (*14*).

### Ion permeation pathway

The central ion-conduction pore contains multiple constrictions that would prevent ion passage (Fig. 4, A and B), as indicated by the pore radius calculation (*35*). Thus, the structure represents a non-conductive conformation, which is consistent with the high pH buffer condition (pH 8.0) used for cryo-EM structure determination. The intimate assembly at the top of the ECD places side chains of F238 in close proximity, generating a constriction that separates the upper and central vestibules. The voluminous and elongated central vestibule is accessible laterally owing to the lack of protein-protein contacts in the middle of the ECD (Fig. 4C). Therefore, the narrow point at F238 might not interfere with ion conduction and could be maintained during the channel gating cycle as ions pass through the three side portals. Further, the slightly positive electrostatic potential of the interior walls of the central vestibule and side entryways would facilitate attraction of extracellular Cl^−^ (Fig. 4C).

**Figure 4.**
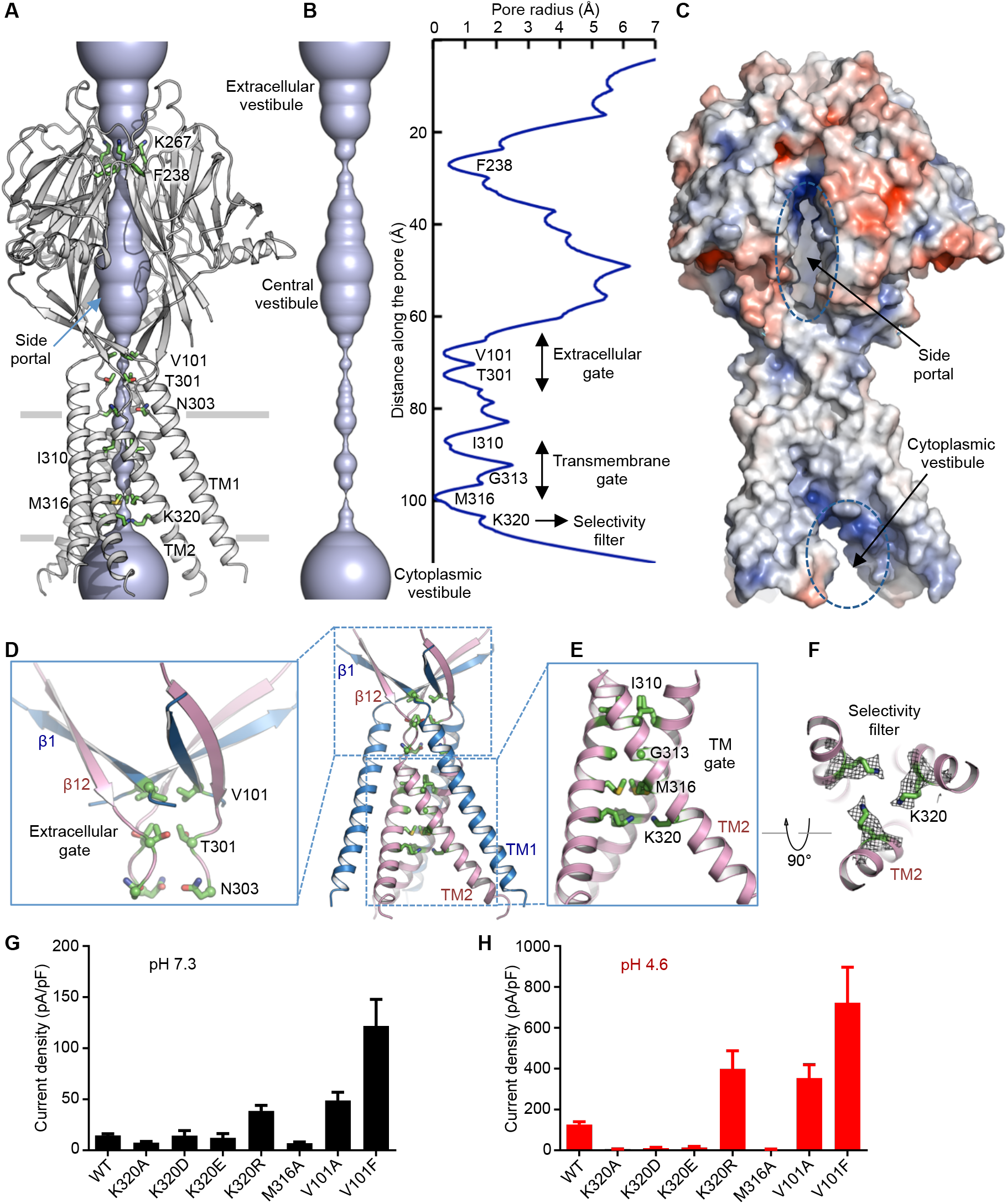
Ion permeation pathway. (**A**) Structure of TMEM206_EM_ and the central ion conduction pore, shown in surface representation. Residues generating constrictions are highlighted and labeled. The side portal is indicated. (**B**) The central ion conduction pore and estimation of the radius (right panel). (**C**) Surface representation of the channel, colored by surface electrostatic potential (red, −5 kT/e; white, neutral; blue, +5 kT/e). The side portal and cytoplasmic vestibule are indicated. (**D**) The extracellular gate at the ECD-TMD junction. V101, T301 and N303 are shown in stick representation. (**E**) The transmembrane gate, constituted by I310, G313 and M316. (**F**) The putative selectivity filter defined by K320. Also shown are side chain densities for K320, contoured at 6.5 σ. (**G-H**) Current densities with an extracellular pH of 7.3 (**G**) and 4.6 (**H**) at +100 mV for TMEM206 mutants. The whole-cell membrane currents were recorded by using voltage ramping from −100 to +100 mV for 500 ms at a holding potential of 0 mV.

At the ECD-TMD junction, the three β1 strands, connected to the outer helices, cross each other and the three β12 strands move inward to join the inner helices (Fig. 4D). This arrangement generates an extracellular gate, which is constituted by V101 in β1 and T301 and N303 in the β12-TM2 linker, above the lipid bilayer (Fig. 4D). Underneath the extracellular gate and within the membrane, the ion-conduction pore is lined by residues from the inner helices TM2, which cross each other at a conserved glycine residue G313 by an angle of ~60° (Fig. 4E). Consecutive constrictions at the pore-facing positions, I310, G313 and M316, appear to form a hydrophobic gate that prohibits ion conduction (Fig. 4E). Beneath the gate, side chains of a highly conserved basic residue K320 point toward the central pore, presumably constituting the anion selectivity filter (Fig. 4F, fig. S1). Immediately below the filter, the ion pore widens substantially on the cytoplasmic side as the inner helices become further apart.

To corroborate our structural findings, we performed mutagenesis studies on key pore-lining residues. The critical role of K320 in channel function was confirmed as substitution of this basic residue with alanine or acidic residues abolished acid-activated *I*_Cl,H_ currents (Fig. 4, G and H, fig. S9, A-C). In marked contrast, the arginine substitution retained channel function (Fig. 4, G and H, fig. S9D), further supporting the requirement of positive charges in the anion-selective filter. Interestingly, M316A also abolished *I*_Cl,H_ currents, suggesting the importance of this pore-facing position right above the selectivity filter (Fig. 4, G and H, fig. S9E). In contrast, the V101A and V101F mutations resulted in increased currents at both pH 7.3 and pH 4.6 (Fig. 4, G and H, fig. S9, F and G), suggesting that V101 is critical to the formation of the extracellular gate.

Previous cysteine-scanning accessibility studies of the entire TM1 and TM2 helices of human TMEM206 revealed that the outer helix TM1 was mostly insensitive to cysteine substitutions (*14*). By contrast, the inner helix TM2 carried multiple cysteine substitutions that displayed either increased or decreased *I*_Cl,H_ currents in response to MTSES application (*14*). For a total of 44 cysteine substitutions, only L309C and K319C in TM2 (corresponding to I310 and K320 in pufferfish TMEM206), failed to elicit acid-activated *I*_Cl,H_ currents (*14*). In light of our structure, these experiments underscore the essential role of these two pore-lining residues. Specifically, I310 is a critical component of the transmembrane gate and K320 forms the anion selectivity filter (Fig. 4, E and F). G313 and M316 are the remaining two residues that constitute the transmembrane gate in pufferfish TMEM206. Interestingly, the corresponding cysteine substitutions in human TMEM206 (G312C and L315C) showed a pronounced MTSES-dependent increase of *I*_Cl,H_ currents (*14*). In addition, the introduction of an acidic residue at position 315 (L315D), which is one helical turn above the anion selectivity filter that is composed of lysine residues (Fig. 4E), rendered the channel nonselective with permeability to both cations and anions (*14*). Collectively, key pore-lining residues with functional implications in gating and selectivity identified by the systematic cysteine substitution experiments are in accordance with our structural and electrophysiological findings. Further, these data support structural conservation between the human and pufferfish orthologs and that our TMEM206_EM_ structure represents a physiologically relevant model for the entire family of Cl^−^ channels.

### Structural convergence of cation and anion channels

Surprisingly, the topology, structure, and assembly of TMEM206 are reminiscent of those of the epithelial sodium channel/degenerin (ENaC/DEG) superfamily of ion channels, including acid-sensing ion channels (ASICs) and epithelial sodium channels (ENaCs) (Fig. 5, A-D), despite a lack of discernible amino acid sequence homology (*36*–*38*). These trimeric channels, selective for oppositely charged ions Na^+^ and Cl^−^, share a common core structure composed of a twelve-stranded β-domain flanked by two transmembrane helices. TMEM206 contains a simple HTH insertion to the outer β-domain, whereas ASICs and ENaCs are furnished with more complex structural elements surrounding the outer β-domain (Fig. 5, A-D). Structures of ASIC1a reveal a central ion permeation path analogous to that of TMEM206, including the upper, central, and cytoplasmic vestibules and multiple constrictions (Fig. 5E and 5F). Additionally, ASIC1a contains an extracellular vestibule and fenestrations, which enlarge upon channel activation to allow ion conduction, leaving the extracellular constrictions along the central path largely unaltered (Fig. 5F) (*36*, *37*). By contrast, in TMEM206, the central vestibule opens to the side and Cl^−^ ions may access the central pore through the three lateral portals (Fig. 5E). However, TMEM206 lacks the extracellular vestibule and fenestrations at the ECD-membrane boundary. Thus, in TMEM206, the extracellular gate between the central vestibule and transmembrane pore needs to expand to pass ions, unless extracellular fenestrations, like those in ASIC1a, are newly synthesized upon channel activation.

**Figure 5.**
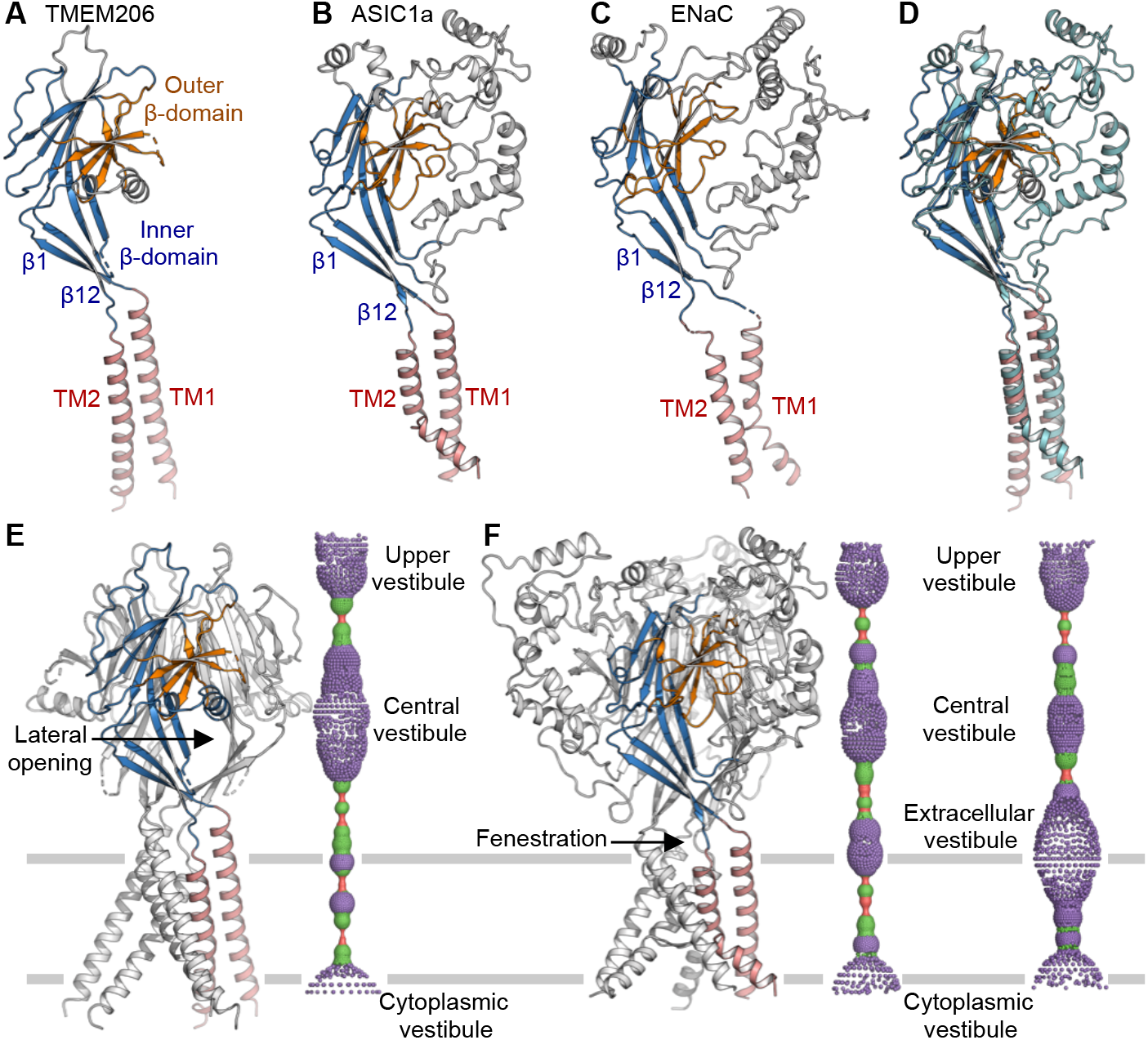
Structural comparison with ASIC and ENaC. (**A**-**C**) Subunit structures of TMEM206 (**A**), ASIC1a (PDB: 6AVE) (**B**), ENaC (PDB: 6BQN) (**C**). Domains are similarly colored. (**D**) Superposition of TMEM206, colored as in (**A**), and ASIC1a colored in cyan. (**E**) The trimeric TMEM206 channel and its central ion conduction pore. The pore is estimated using the program HOLE and depicted as colored dots (pore radius: red < 1.15 Å < green < 2.3 Å < blue). (**F**) The trimeric ASIC1a channel and the central pore in the closed (middle panel, PDB: 6AVE) and open (right panel, PDB: 4NTW) states.

ENaCs are activated by release of inhibitory peptides through proteolysis in the ECD, whereas ASICs and TMEM206 are activated by extracellular protons (*13*, *14*, *37*, *38*). In ASIC1a, an electrostatically negative ‘acidic pocket’, formed between the β-domain and peripheral domains at the subunit interface, adopts an expanded conformation in the resting state and collapses upon exposure to extracellular protons, resulting in expansion of the lower β-domain and an iris-like opening of the transmembrane gate (*36*, *37*). In contrast, TMEM206 lacks the additional structural elements involved in formation of the ‘acidic pocket’ as in ASIC1a, suggesting a distinct acid-sensing mechanism.

Nevertheless, conserved structural features between TMEM206 and ASICs suggest the possibility of analogous gating conformational changes, such as a relatively stationary structural scaffold at the upper ECD, expansion of the lower β-domain, and an iris-like opening of the gate in the membrane.

## Discussion

Single-particle cryo-EM has facilitated structure determination of integral membrane proteins that are unattainable using traditional X-ray crystallography. However, achieving near-atomic resolution for membrane proteins of small size remains a major technical challenge due to low contrast- and signal-to-noise ratios (*39*). In this study, we obtained a 3.5 Å resolution structure of a channel with an ordered portion of only ~90 kDa. This was made possible by fusion with a small crystallization chaperone BRIL, which has proven to be useful in improving stability of otherwise sub-optimal membrane proteins and in promoting crystal packing (*33*). The improvement in resolution was enabled by markedly increased particle density on the cryo-EM grids as a result of enhanced homogeneity and reduced aggregation, rather than a gain of molecular mass as BRIL was invisible in the density map. Perhaps owing to the flexible connection between the channel and BRIL, the structural and functional integrity is maintained in the fusion construct TEME206_EM_. Analogous to crystallography, the BRIL fusion strategy may be broadly applicable to single-particle cryo-EM studies of small-sized and intrinsically unstable membrane proteins.

This work now defines the structure and assembly of a new class of Cl^−^ channels TMEM206 and establishes a framework for further functional and mechanistic investigation. To provide structural insights into proton activation, we attempted to determine structures of TMEM206_EM_ at low pH (~5.0) but were unable to obtain meaningful cryo-EM reconstructions. Strikingly, the structure of the TMEM206 Cl^−^ channel closely resembles those of Na^+^-selective channels ASICs and ENaCs, which are unrelated in amino acid sequence and conduct positively, instead of negatively, charged ions (*36*–*38*). The conserved core structure suggests that these channels, selective for either cations or anions, may experience similar gating conformational changes.

## Materials and Methods

### Protein expression and purification

The codon-optimized DNA fragments encoding sixteen TMEM206 orthologs were synthesized (Bio Basic Inc). For overexpression in yeast *Pichia pastoris*, DNA fragments were transferred into a modified pPICZ-B vector with a PreScission protease cleavage site followed by a C-terminal GFP-His_10_ tag. The pufferfish TMEM206 was identified as a promising candidate for structural studies, as evaluated by fluorescence-detection size-exclusion chromatography (FSEC). Initial cryo-EM analysis using the full-length wild-type protein showed that purified channels were prone to aggregation on cryo-EM grids, precluding structure determination to high resolution. To improve the biochemical stability of the channel, BRIL (thermo-stabilized apocytochrome b_562_RIL) was fused to the C-terminus of pufferfish TMEM206. To increase the structural rigidity of the fusion construct, the last four C-terminal residues of TMEM206 were further removed. The expression construct TMEM206_EM_ includes residues 1-349 of pufferfish TMEM206 and a C-terminal BRIL followed by the PreScission protease cleavage site and GFP-His_10_ tag. For electrophysiological recordings, DNA fragments were ligated into a modified pCEU vector containing a C-terminal GFP-His_8_ tag. Mutations used in this study were generated by site-directed mutagenesis.

Yeast cells expressing the full-length wild-type pufferfish TMEM206 were disrupted by milling (Retsch MM400), and resuspended in buffer containing 50 mM Tris pH 8.0 and 150 mM NaCl supplemented with DNase I and protease inhibitors including 2.5 μg/ml leupeptin, 1 μg/ml pepstatin A, 100 μg/ml 4-(2-Aminoethyl) benzenesulfonyl fluoride hydrochloride, 3 μg/ml aprotinin, 1 mM benzamidine and 200 μM phenylmethane sulphonylfluoride. The cell mixture was extracted with 1% (w/v) Lauryl Maltose Neopentyl Glycol (LMNG, Anatrace) for 2 h with stirring at 4°C, and then centrifuged for 1 h at 30,000 g. The supernatant was collected and incubated with 3 ml of cobalt-charged resin (G-Biosciences) for 3 h at 4°C. Resin was then collected and washed with 30 ml of buffer 20 mM Tris pH 8.0, 150 mM NaCl, 20 mM imidazole, and 85 μM glyco-diosgenin (GDN, Anatrace). The channel protein was eluted with 200 mM imidazole and digested with PreScission protease at 4°C overnight to remove the C-terminal GFP-His_10_ tag. Further purification was performed on a Superose 6 Increase 10/300 gel filtration column (GE Healthcare Life Sciences) in 20 mM Tris pH 8.0, 150 mM NaCl, and 40 μM GDN. Peak fractions containing channel protein were concentrated to ~6 mg/ml and used immediately for cryo-EM grid preparations. Purification of TMEM206_EM_ followed the same procedure as that for the wild type, except that 0.05 mg/ml Soybean polar lipid extract (Avanti Polar Lipids, Inc.) was included in the wash, elution and gel filtration buffers.

### Cryo-EM grid preparation and imaging

Cryo-EM grids were prepared by applying freshly purified channel protein (~3.5 μl) to glow-discharged copper Quantifoil R2/2 holey carbon grids (Quantifoil), which were then blotted for 2 s with ~100% humidity and flash frozen in liquid ethane using FEI Vitrobot Mark IV (FEI). Images were collected using a Titan Krios (FEI) electron microscope operating at 300 kV with a Gatan K2 Summit (Gatan) detector and GIF Quantum energy filter with a slit width of 20 eV. Data collection was performed using EPU software (https://www.fei.com/software/epu-automated-single-particles-software-for-life-sciences/) in the super-resolution mode with a pixel size of 0.55 Å and a nominal defocus range of −1.0 to −2.5 μm. With a dose of ~7.8 electrons per Å^2^ per second, each micrograph was recorded for 8 s in 40 frames (an accumulated dose of ~62 electrons per Å^2^).

### Data processing and map calculation

Recorded micrographs were motion-corrected and dose-weighted using MotionCor2 (*40*). Does-weighted micrographs were subjected to contrast transfer function (CTF) determination using GCTF (*41*). Following motion correction and CTF estimation, micrographs of poor quality were manually removed from the datasets. For the pufferfish TMEM206_EM_ dataset, LoG-based auto-picking was used to pick 7629 particles from randomly selected micrographs to generate 2D classes for automatic picking in RELION3 (*42*). Good 2D classes (2827 particles) were used as templates for auto-picking from 3,411 micrographs, resulting in a total of 1,809,512 particles. Particles were extracted using a box size of 220 pixels and subjected to 2D classification with a mask diameter of 170 Å. After two rounds of 2D classification, 835,518 particles were selected and imported into cryoSPARC (*43*). An initial map was then generated in cryoSPARC and used for 3D auto-refinement in RELION3, resulting in a 4.03 Å resolution map. Further 3D classification requesting six classes identified two classes showing complete channel features (505,115 particles). These two classes were combined and subjected to 3D refinement followed by CTF refinement and Bayesian polishing. The final reconstruction achieved an overall resolution of 3.46 Å from masked refinement with C3 symmetry.

For the full-length wild-type pufferfish channel, automatic picking resulted in 430,043 particles from 1,308 micrographs. After two rounds of 2D classification, 49,851 particles were selected to create an initial map in cryoSPARC for 3D classification requesting four classes in RELION3. Two classes corresponding to a complete channel (31,362 particles) were selected and subjected to masked 3D refinement, yielding a reconstruction with an overall resolution of 6.17 Å after post-processing.

### Model building, refinement and validation

*De novo* model building, guided by densities for bulky side chains and disulfide bonds, was conducted in COOT (*44*). Cycles of model building in COOT and real space refinement using real_space_refine against the full map in PHENIX (*45*) were performed to obtain the final refined atomic model, which was validated using MolProbity (*46*). The final model includes residues 65-159, 168-251, and 255-334. The densities for side chains of residues 65-76 and 321-334 are poorly defined, and therefore these residues were modeled as alanine. Pore radius calculation was performed using the program HOLE (*35*). Structural figures were rendered using PyMol (pymol.org).

### Electrophysiology

Knockout of the human *TMEM206* gene in the HEK293T cell line was conducted using CRISPR/Cas9-mediated gene disruption (*47*). The *TMEM206* gene was targeted using the reported guideRNA (gRNA, 5’-GGACCGAGAAGACGTTCTTC-3’, negative strand) sequence (*13*). The gRNA was inserted into PX459 V2.0 plasmid (Addgene, catalog #62988) and transfected into HEK293T cells using FuGENE HD transfection reagent (Promega). After 24 h, cells were transferred to fresh medium with 5 μg/ml puromycin for additional 7 days. Single colonies were then isolated using limiting dilution. The knockout cell line was determined through genotyping analysis of frameshift mutations by target-site-specific PCR and cloning followed by Sanger sequencing.

The *TMEM206* knockout cell line was used for electrophysiology experiments. The C-terminal GFP-tagged wild-type pufferfish TMEM206 channel or each mutant channel was transfected. Whole-cell patch clamp recordings were performed at room temperature using an Axon 700B amplifier (Molecular Devices). Pipettes were pulled from borosilicate glass (BF 150-86-10; Sutter Instrument) with a Sutter P-1000 pipette and filled with the intracellular solution containing 135 mM CsCl, 1 mM MgCl_2_, 2 mM CaCl_2_, 10 mM HEPES, 5 mM EGTA, 4 mM MgATP (280-290 mOsm/kg; pH 7.2 with CsOH). The external solution contained 145 mM NaCl, 2 mM KCl, 2 mM MgCl_2_, 1.5 mM CaCl_2_, 10 mM HEPES, 10 mM glucose (300 mOsm/kg; pH 7.3 with NaOH). To make different acidic pH solutions, 5 mM Na_3_-citrate was used as buffer and the pH was adjusted using citric acid. Holding at 0 mV, voltage ramp from −100 to +100 mV for 500 ms was used to record whole-cell currents. For anionic selectivity experiments, bath solutions were replaced with 145 mM NaX/5mM Na_3_-citrate (pH 4.6 was adjusted with citric acid), where X was Cl^−^, Br^−^ and I^−^, respectively. Data were acquired using Clampex 10.4 software (Molecular Devices). Currents were filtered at 2 kHz and digitized at 10 kHz. Data were analyzed and plotted using Clampfit 10 (Molecular Devices). The concentration–response curve was fitted with the logistic equation: Y = Y_min_ + (Y_max_−Y_min_)/(1 + 10^[(logpH_50_−X) × Hill slope]), where Y is the response at a given pH, Y_max_ and Y_min_ are the maximum and minimum responses, X is the logarithmic value of the pH and Hill slope is the slope factor of the curve. pH_50_ is the pH value that gives a response halfway between Y_max_ and Y_min_. The permeability ratios were calculated from shifts in reversal potential with the Goldman-Hodgkin-Katz Equation. All data are presented as mean ± SEM.

## Acknowledgments

We thank members of the Yuan laboratory for discussion.

## Author contributions

Z.D. designed the experiments, performed biochemical preparations, cryo-EM experiments, structural determination and analysis. J.Z. aided biochemical preparations. Y.Z., J.F. and H.H conducted electrophysiology experiments. H.Z. generated the KO cell line for electrophysiology. M.R., J.A.J.F., and Z.D. performed cryo-EM data acquisition. P.Y. conceived and supervised the project. Z.D., Y.Z., J.F., H.H, and P.Y. analyzed the results and prepared the manuscript with input from all authors.

## Data and materials availability

The cryo-EM maps have been deposited to Electron Microscopy Data Bank with accession codes EMD-22342 and EMD-22343. Atomic coordinates have been deposited to the Protein Data Bank (PDB) with accession code 7JI3. Correspondence and requests for materials should be addressed to P.Y.

---

**Fig S1.**
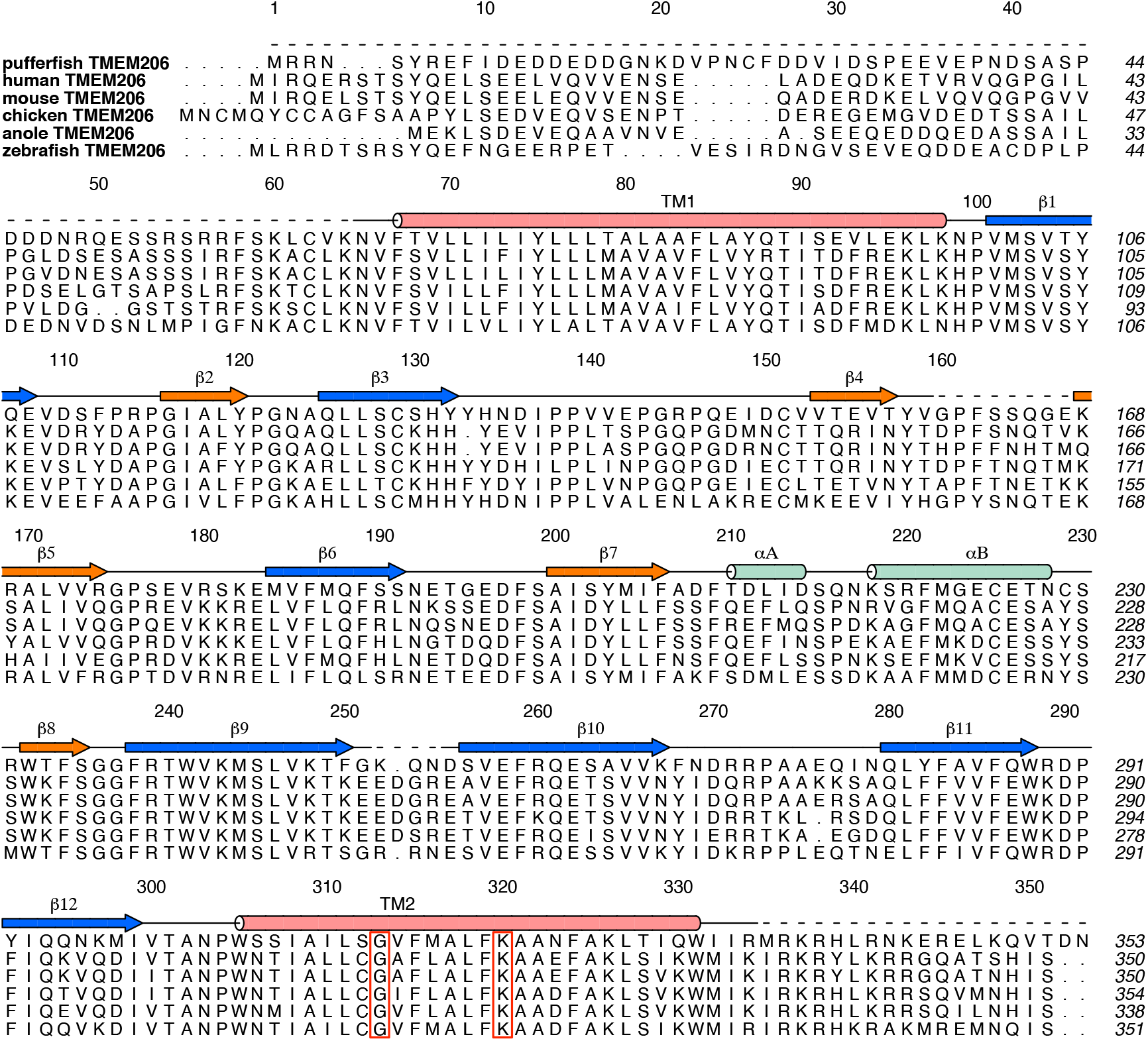
Sequence alignment of TMEM206 orthologs. The protein sequences of pufferfish (NCBI sequence: XP_011618135.1), human (NCBI sequence: NP_060722.2), mouse (NCBI sequence: NP_080140.1), chicken (NCBI sequence: XP_015139352.1), anole (NCBI sequence: XP_003216059.1), and zebrafish (NCBI sequence: NP_001278691.1) TMEM206 were aligned. Secondary structure elements of pufferfish TMEM206 are also shown. Dashed lines indicate unresolved regions in the cryo-EM density. Conserved residues G313 and K320 are highlighted in red boxes.

**Fig S2.**
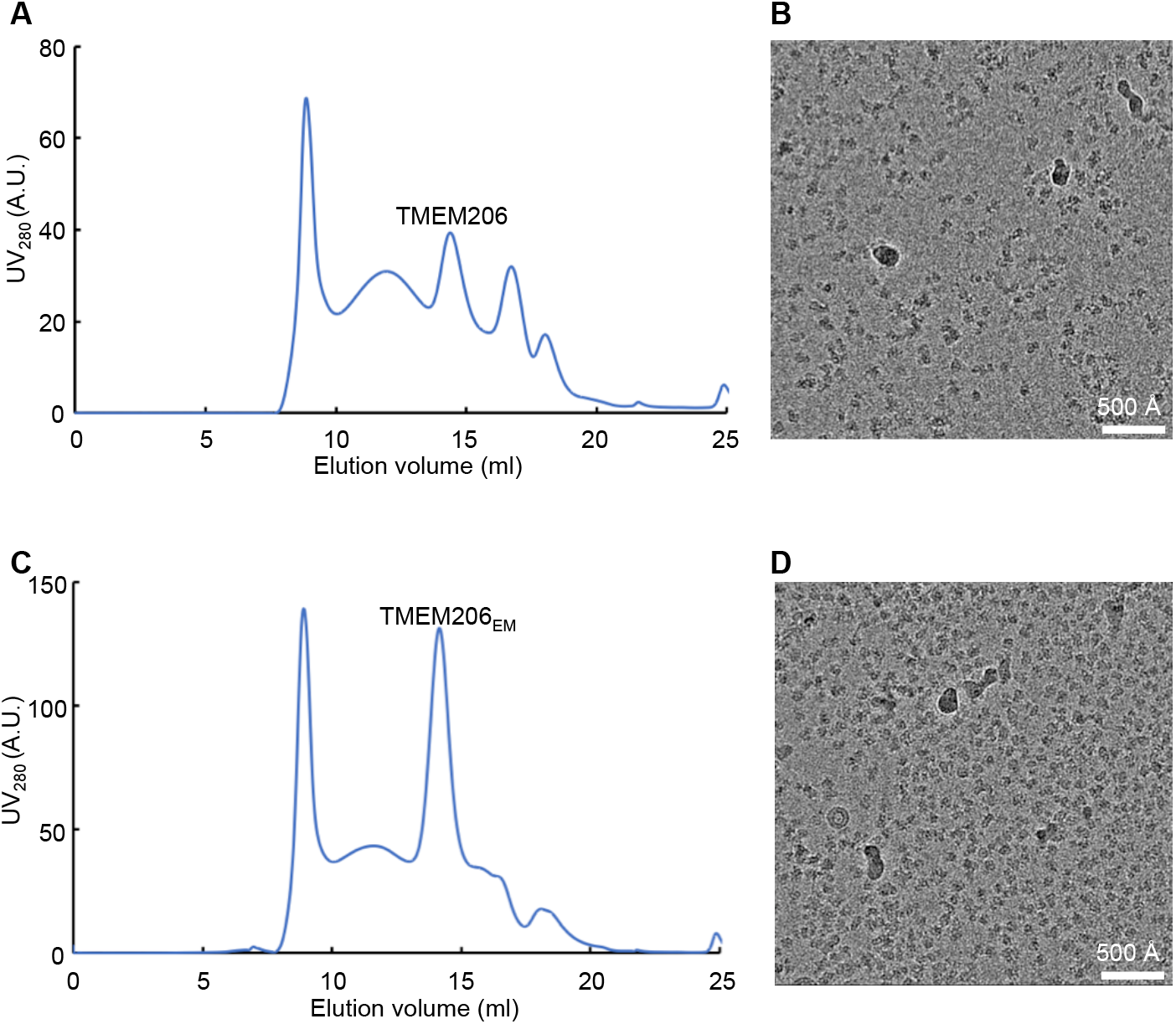
BRIL fusion increases particle density. (**A**) Purification of the wild-type full-length pufferfish TMEM206 channel on size-exclusion chromatography. The oligomeric channel is indicated. (**B**) Representative cryo-EM micrograph of pufferfish TMEM206. (**C**) Purification of the channel-BRIL fusion construct TMEM206_EM_. (**D**) Representative cryo-EM micrograph of TMEM206_EM_.

**Fig S3.**
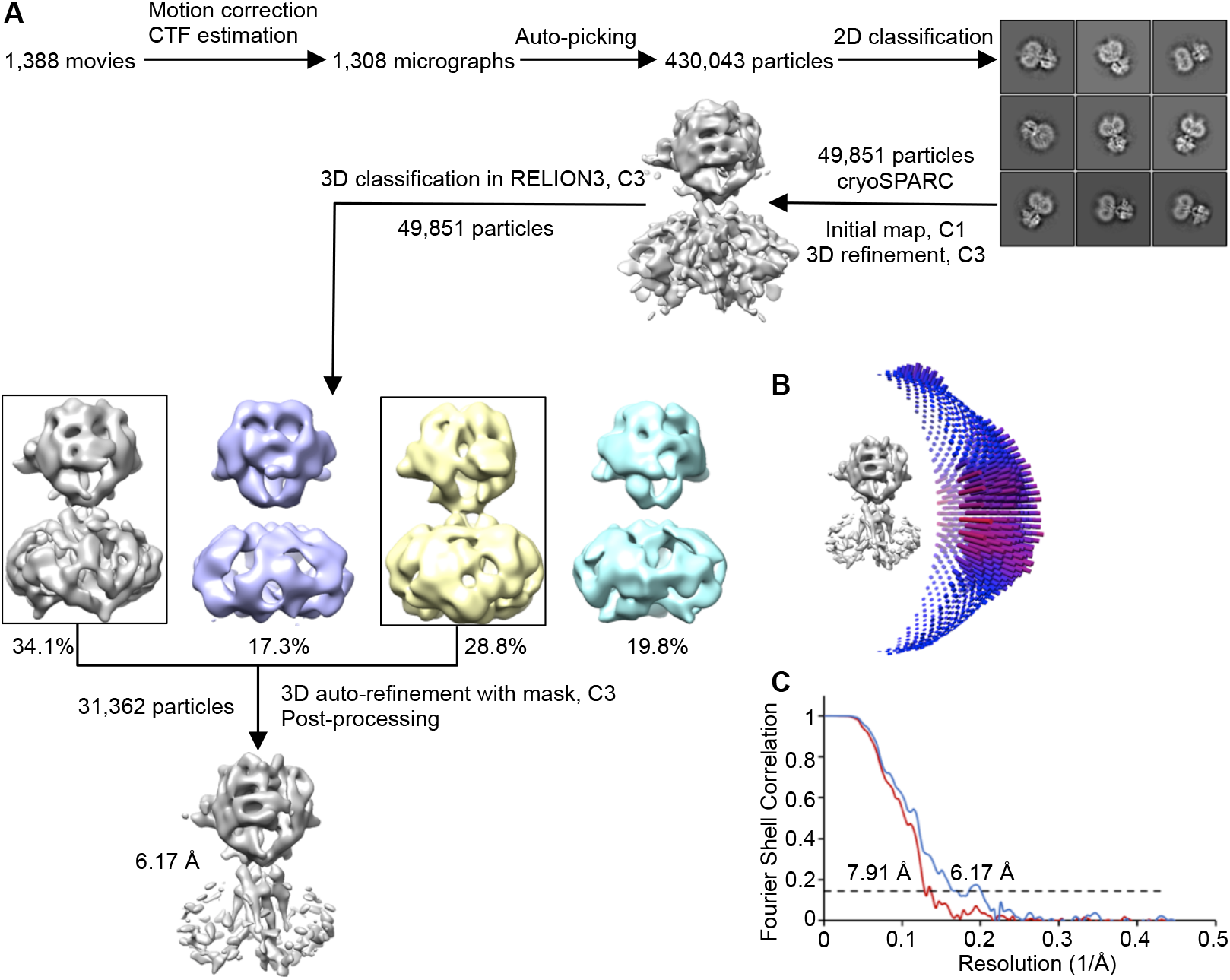
Cryo-EM reconstruction of pufferfish TMEM206. (**A**) Schematic of cryo-EM data processing. (**B**) Euler distribution of particles included in the final reconstruction. C3 symmetry was imposed. (**C**) Fourier shell correlation before and after post-processing in RELION3.

**Fig S4.**
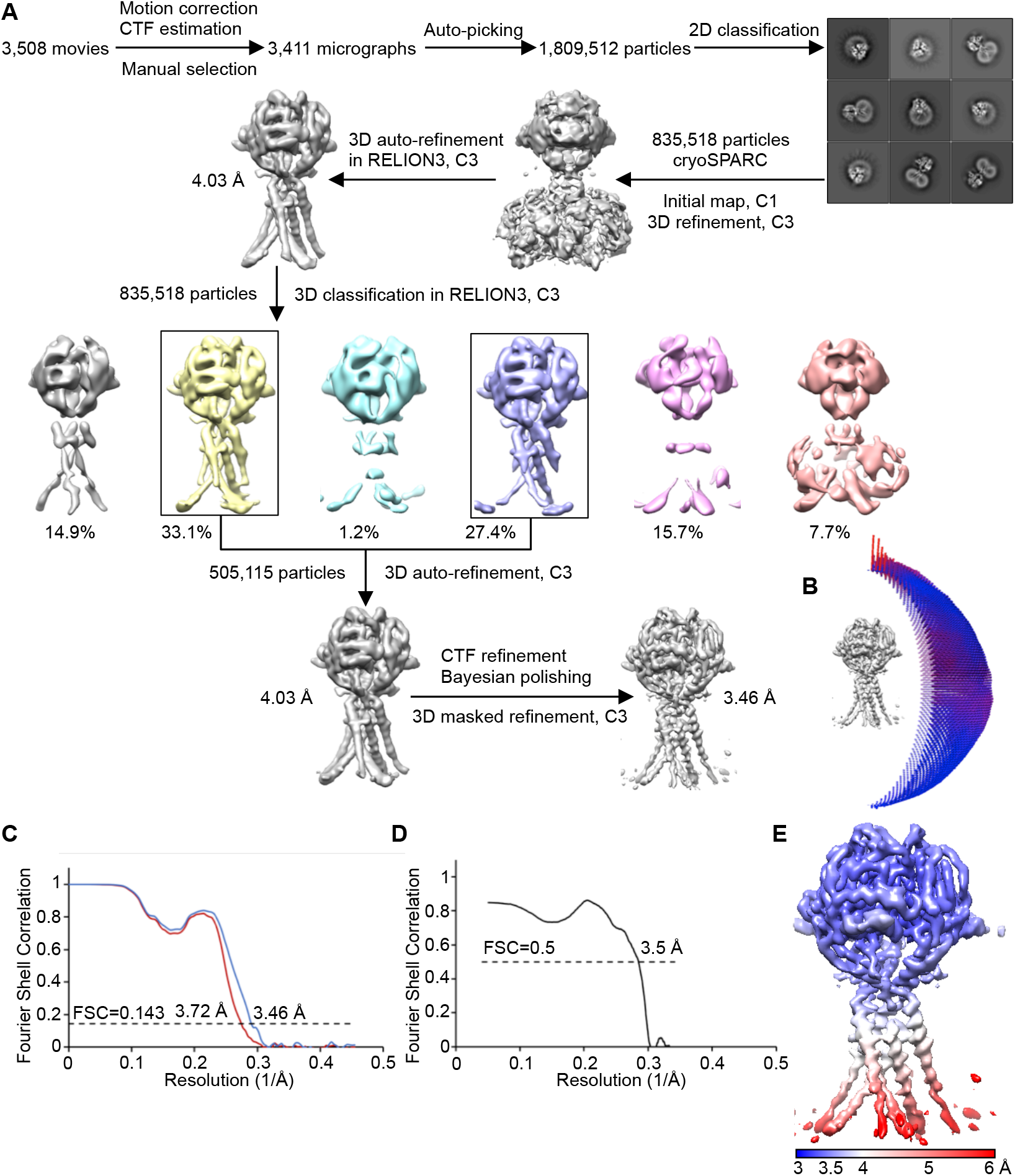
Cryo-EM reconstruction of TMEM206_EM_. (**A**) Schematic of cryo-EM data processing. (**B**) Euler distribution of particles included in the final reconstruction. C3 symmetry was imposed. (**C**) Fourier shell correlation for unmasked (red) and masked (blue) refinement in RELION3. (**D**) Fourier shell correlation between the refined model and the full map. (**E**) Cryo-EM density map colored by local resolution.

**Fig S5.**
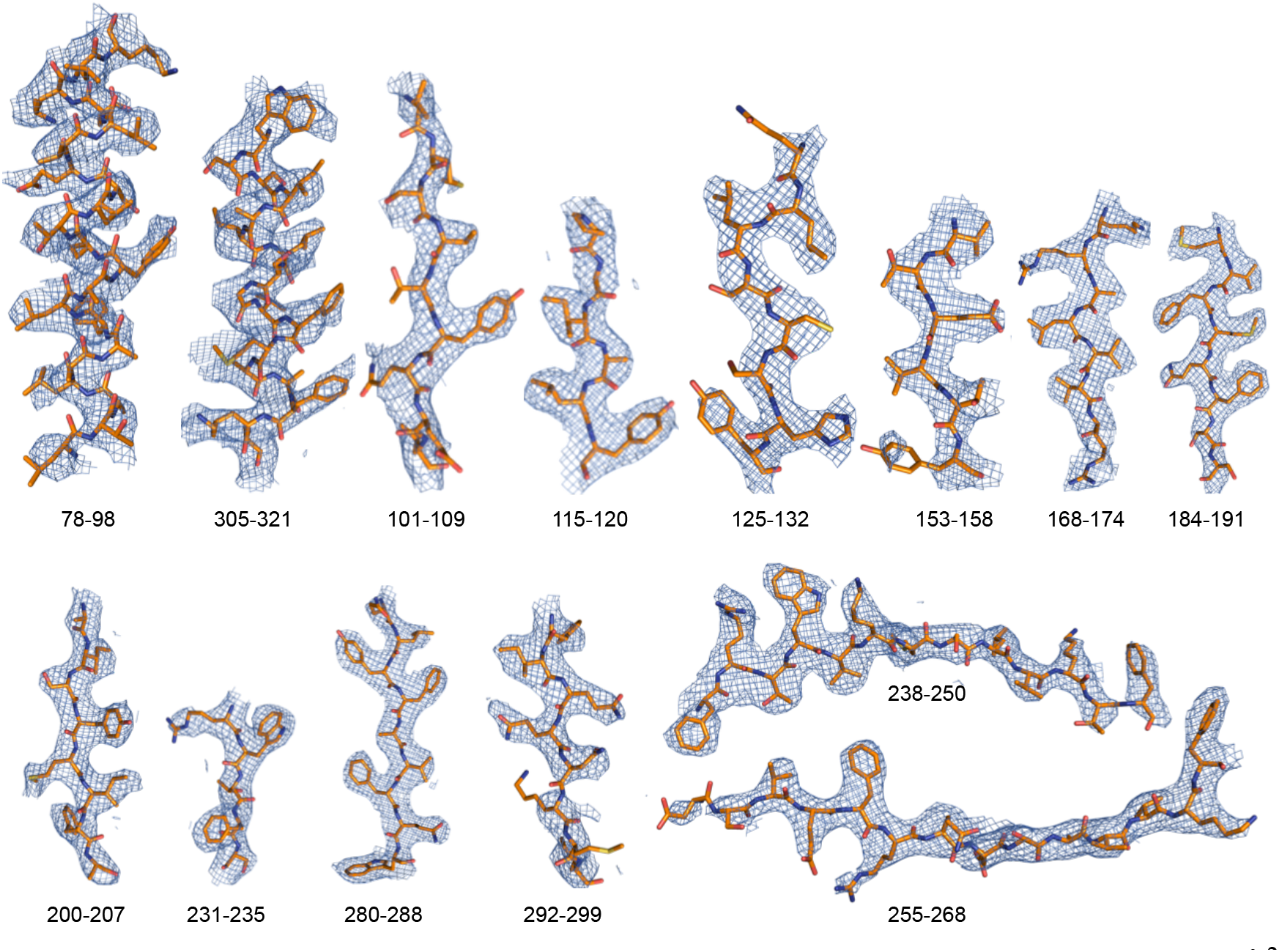
Cryo-EM densities. The map was sharpened by applying a B-factor of −147 Å^2^ in RELION3.

**Fig S6.**
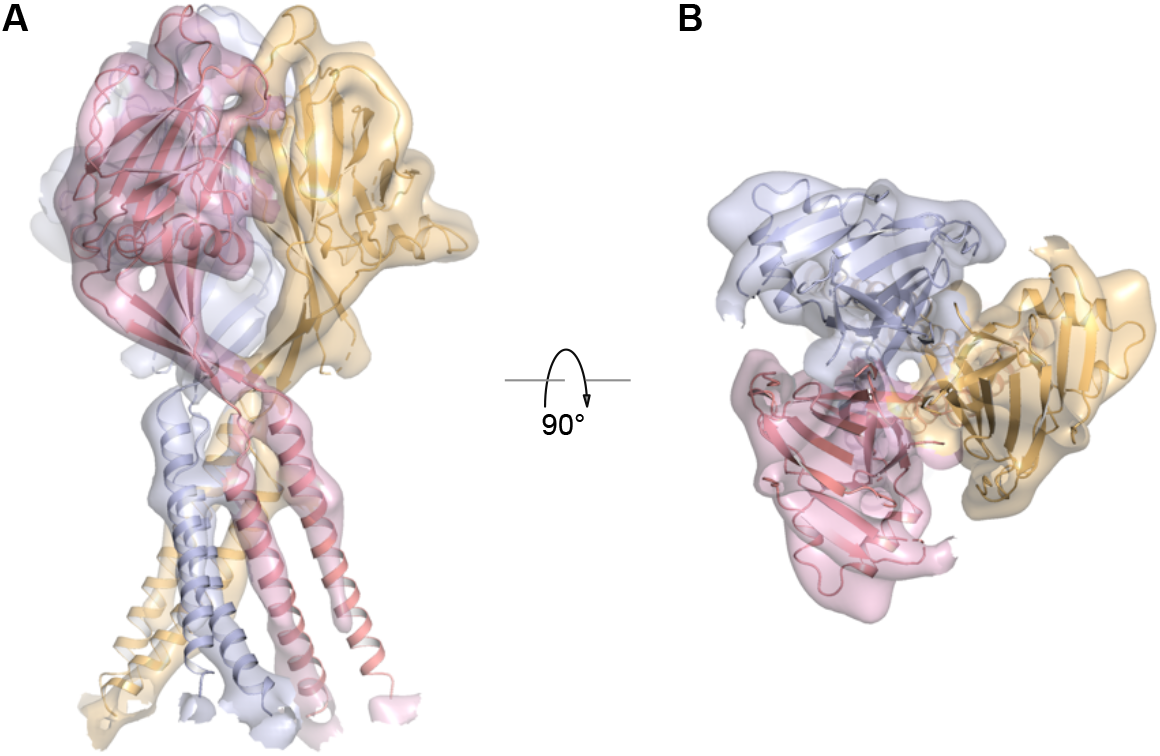
TMEM206_EM_ structure matches that of the wild-type full-length channel. (**A**-**B**) Orthogonal views of the TMEM206_EM_ structure fitted into the low-resolution ryo-EM density (contoured at 7.0 σ) of the intact wild-type channel.

**Fig S7.**
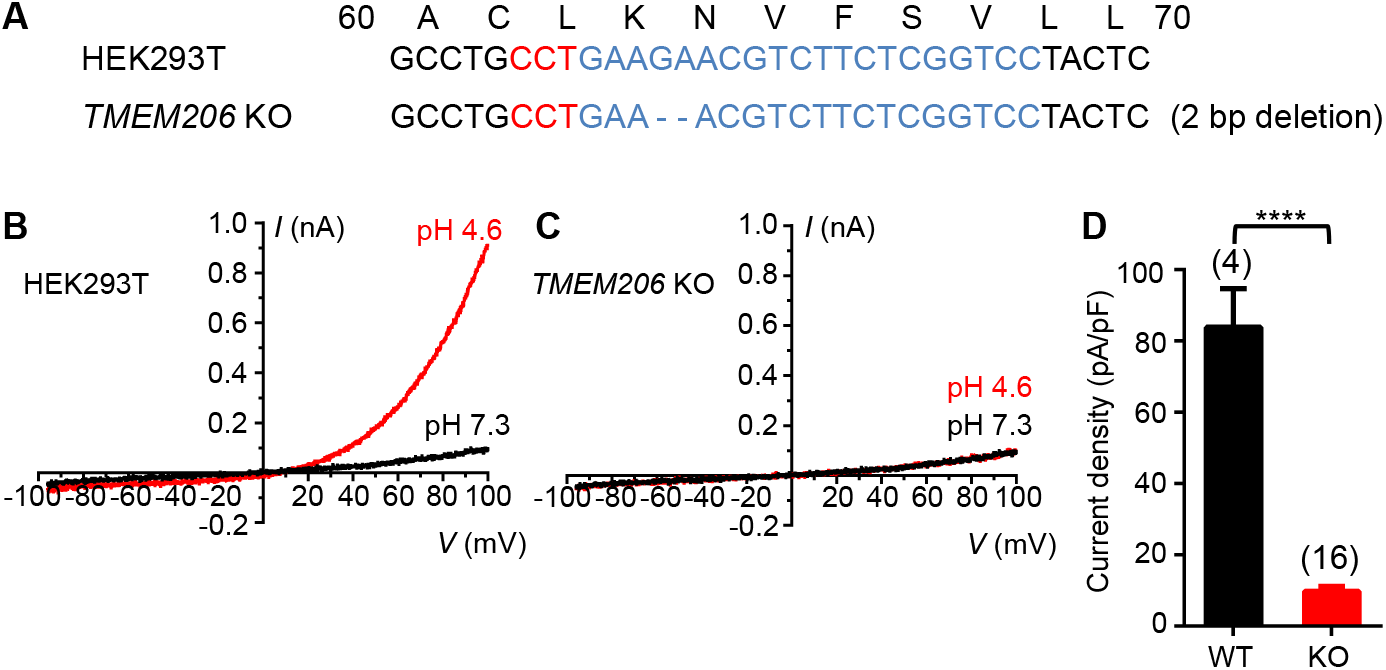
Endogenous human TMEM206-mediated *I*_Cl,H_ current is essentially abolished in the *TMEM206* knockout (KO) HEK293T cell line. (**A**) Knockout (KO) of the *TMEM206* gene in HEK293T using CRISPR. Protospacer adjacent motif (PAM) and targeted DNA sequences are highlighted in red and blue, respectively. The *TMEM206* gene was inactivated by a 2-bp deletion in the KO cell line. (**B**-**C**) Whole-cell currents induced by extracellular pH 4.6 in wild-type (WT) (**B**) and *TMEM206* KO (**C**) HEK293T cells. (**D**) Comparison of acid-induced current densities at pH 4.6 between WT and *TMEM206* KO HEK293T cells. Data are showed as mean ± SEM. ****p < 0.0001, Student’s t-test.

**Fig S8.**
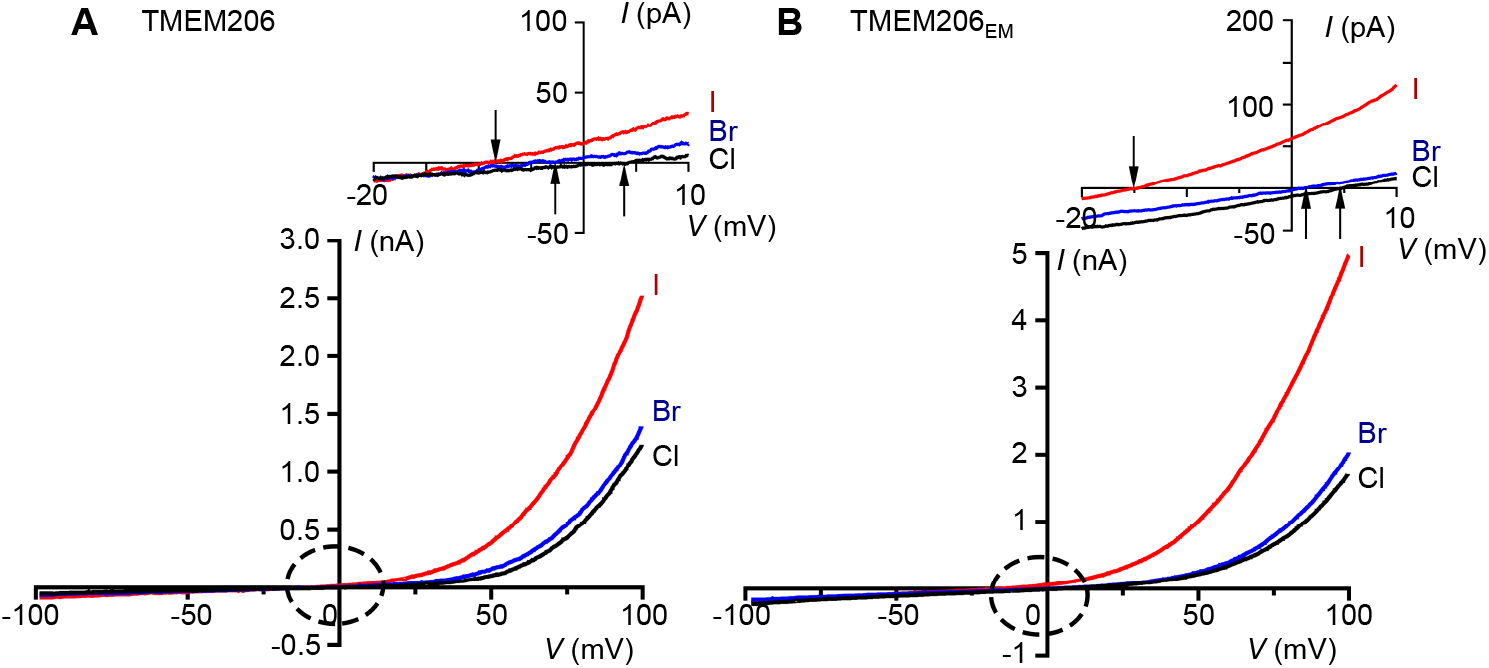
Comparable anion selectivity between pufferfish TMEM206 and TMEM206_EM_. (**A**-**B**) Representative I–V relationships (at pH 4.6) recorded with extracellular solutions containing I^−^, Br^−^, or Cl^−^ for pufferfish TMEM206 (**A**) and TMEM206_EM_ (**B**). Arrows indicate the reversal potentials in the close-up images.

**Fig S9.**
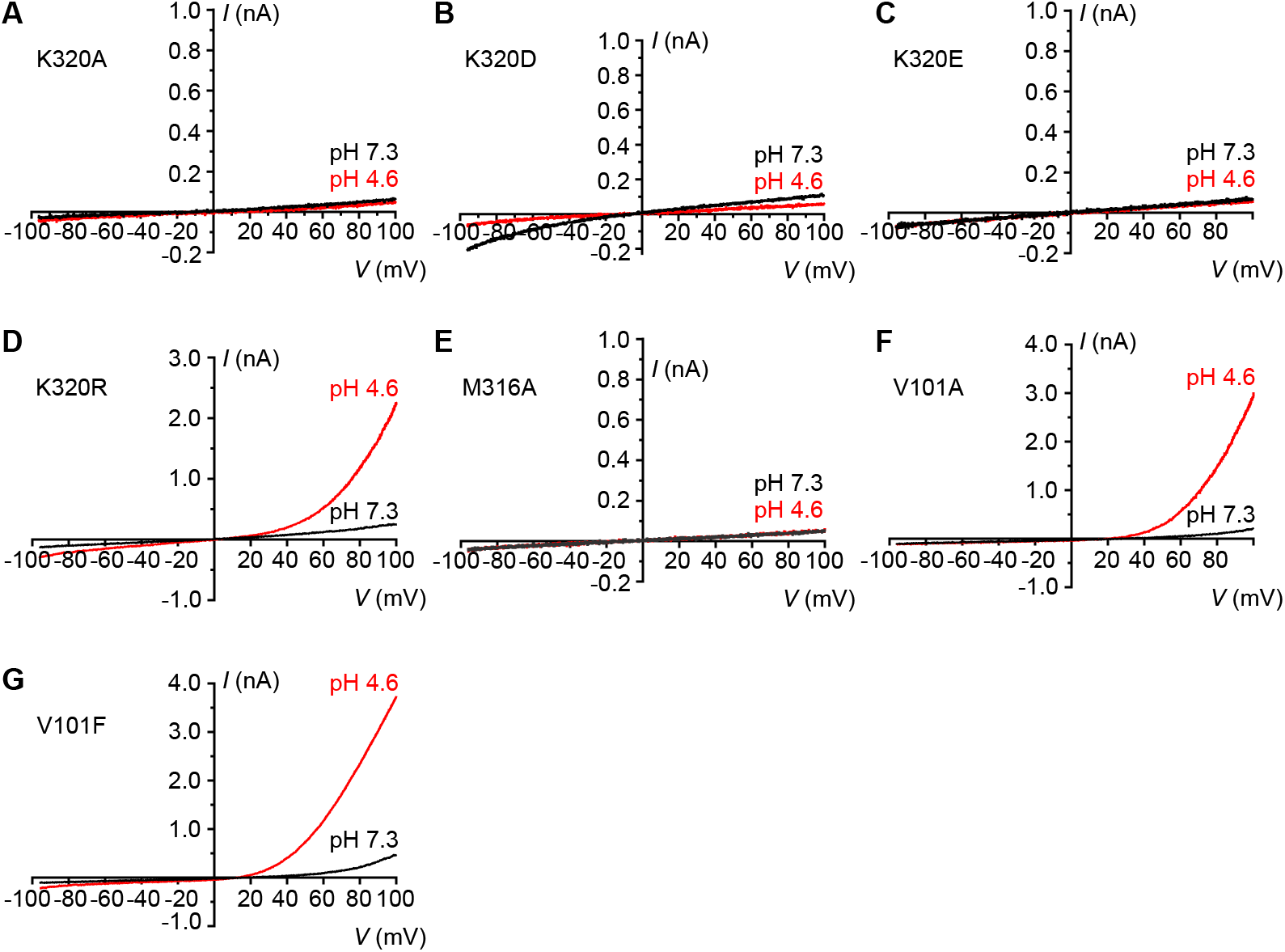
Electrophysiological characterization of pufferfish TMEM206 mutants. (**A**-**G**) Representative ramp current traces of proton-activated chloride currents activated by pH 4.6 for pufferfish TMEM206 mutants K320A (**A**), K320D (**B**), K320E (**C**), K320R (**D**), M316A (**E**), V101A (**F**) and V101F (**G**).

**Table S1.** Cryo-EM data collection, refinement and validation statistics.

|  | TMEM206<sub>EM</sub><br>(EMD-22343)<br>(PDB: 7JI3) | TMEM206<br>(EMD-22342) |
| --- | --- | --- |
| **Data collection and processing** |  |  |
| Magnification | 105,000 | 105,000 |
| Voltage (kV) | 300 | 300 |
| Electron exposure (e-/Å<sup>2</sup>) | 62 | 62 |
| Defocus range (μm) | -1.0 to -2.5 | -1.0 to -2.5 |
| Pixel size (Å) | 1.1 | 1.1 |
| Symmetry imposed | C3 | C3 |
| Initial particle images (no.) | 1,809,512 | 430,043 |
| Final particle images (no.) | 505,115 | 31,362 |
| Map resolution (Å) | 3.46 | 6.17 |
| FSC threshold | 0.143 | 0.143 |
| Map resolution range (Å) | 3.0-6.0 |  |
| **Refinement** |  |  |
| Initial model used (PDB code) | N/A |  |
| Model resolution (Å) | 3.50 |  |
| FSC threshold | 0.5 |  |
| Model resolution range (Å) | 3.30 |  |
| Map sharpening *B* factor (Å<sup>2</sup>) | -147 |  |
| Model composition |  |  |
| Non-hydrogen atoms | 6,000 |  |
| Protein residues | 777 |  |
| Ligands | 0 |  |
| *B* factors (Å<sup>2</sup>) |  |  |
| Protein | 17.2 |  |
| Ligand | N/A |  |
| R.m.s. deviations |  |  |
| Bond lengths (Å) | 0.005 |  |
| Bond angles (°) | 0.785 |  |
| Validation |  |  |
| MolProbity score | 1.61 |  |
| Clashscore | 6.77 |  |
| Poor rotamers (%) | 0 |  |
| Ramachandran plot |  |  |
| Favored (%) | 96.44 |  |
| Allowed (%) | 3.56 |  |
| Disallowed (%) | 0 |  |

